# Pinc: a simple probabilistic AlphaFold interaction score

**DOI:** 10.64898/2026.03.02.708997

**Authors:** Mihaly Badonyi, Agnes Toth-Petroczy

## Abstract

AlphaFold has enabled large-scale prediction of protein–protein and protein–nucleic acid complexes, but ranking and assessing the quality of predicted models remain challenging. Existing confidence scores are often highly parametrised and provide limited interpretability. We introduce a simple geometric framework that converts AlphaFold predicted aligned error (PAE) into conditional contact probability. We show that these probabilities are well calibrated to the fraction of native contacts observed across experimentally determined structures. Motivated by this, we define the Pinc score (**P**robability of **i**nterface **n**ative **c**ontacts) as the mean contact probability between interacting chains. Because the probabilistic interpretation extends to individual residues, Pinc captures local structural constraint beyond interfacial burial, enabling residue-level prioritisation of hotspot positions for mutational studies. Depending solely on a single empirically fixed contact radius, Pinc offers an interpretable path from PAE to interface confidence, matching or exceeding the classification performance of more complex methods across five independent benchmark sets. We provide a portable, dependency-free `C` program and a Google Colab notebook for calculating Pinc scores for AlphaFold models at https://git.mpi-cbg.de/tothpetroczylab/Pinc.

## 1 Introduction

The AlphaFold2-based AlphaFold-Multimer (Evans et al. 2021; Jumper et al. 2021) and the more recent AlphaFold3 (Abramson et al. 2024)—collectively referred to here as AlphaFold—have become the gold standard for modelling the structures of interacting proteins. Central to these approaches is their ability to self-assess predictions by generating scores that quantify the reliability of an interface. Such confidence scores can distinguish biologically relevant interfaces from stochastic associations, and provide a basis for evaluating both individual predictions and large-scale interaction screens, a need reflected in community-wide model assessment efforts such as CASP (Kryshtafovych et al. 2026) and CAPRI (Lensink et al. 2023). However, these scores generally do not lend themselves to a simple probabilistic interpretation.

The primary AlphaFold interaction confidence score is ipTM [interface predicted template modeling (TM) score (Evans et al. 2021; Jumper et al. 2021)]. The TM-score is a measure of structural similarity originally developed to evaluate homology models by providing a chain length-independent score of overall fold similarity (Zhang and Skolnick 2004). AlphaFold uses the predicted aligned error (PAE), which quantifies the expected positional error for each residue pair, to estimate the TM-score of the predicted interface relative to the corresponding region in the (unknown) true structure. Despite its utility, the TM-score, and by extension ipTM, has a nontrivial relationship to interaction probability, as a value of 0.5 is already highly statistically significant under a null model of random structural matches (Xu and Zhang 2010).

While the TM-score was designed to be length-independent, its implementation in ipTM is sensitive to total chain length, particularly when regions outside of the interface are unstructured. These and other limitations have motivated interface-centric metrics, including pDockQ (Bryant et al. 2022), iLIS (Kim et al. 2024, 2026), actifpTM (Varga et al. 2025), and ipSAE (Dunbrack Jr 2025), which aim to reduce the influence of global structure on interaction confidence. For example, pDockQ is a sigmoid-transformed score combining the number of interface contacts and the average pLDDT of interface residues to predict DockQ (Bryant et al. 2022), a composite quality score developed for protein–protein docking models (Basu and Wallner 2016). As a result, pDockQ lacks a direct probabilistic interpretation. Similarly, because actifpTM and ipSAE are formulated relative to the TM-score, and iLIS is derived from PAE and distance cutoffs, interpreting these metrics and their biologically meaningful thresholds can be challenging.

Here, we develop an intuitive interface confidence score using a geometric model that maps PAE values to residue–residue contact probabilities. We show that interchain contact probabilities align closely with the fraction of native contacts in two protein interaction benchmarks. Inspired by this correspondence, we define Pinc (**P**robability of **i**nterface **n**ative **c**ontacts) as the mean interface contact probability, which estimates the fraction of native contacts. Evaluated on five independent benchmark sets spanning diverse interaction classes, Pinc yields performance on par with state-of-the-art metrics, suggesting that more complex methods offer diminishing returns. As the most interpretable direct mapping from PAE to interface confidence currently available, Pinc provides a parsimonious alternative for evaluating AlphaFold predictions.

## 2 Results

### 2.1 Development of the Pinc score

We derive conditional residue–residue contact probabilities from the PAE matrix using a geometric model that assumes positional uncertainty is isotropic, defining a spherical uncertainty region around each residue’s predicted position (Pellegrini et al. 2025). The contact probability for a scored residue is the volume of intersection between its uncertainty sphere and the contact sphere of the aligned residue, normalised by the volume of the former. An overview of the method is shown in Figure 1a.

**Figure 1:**
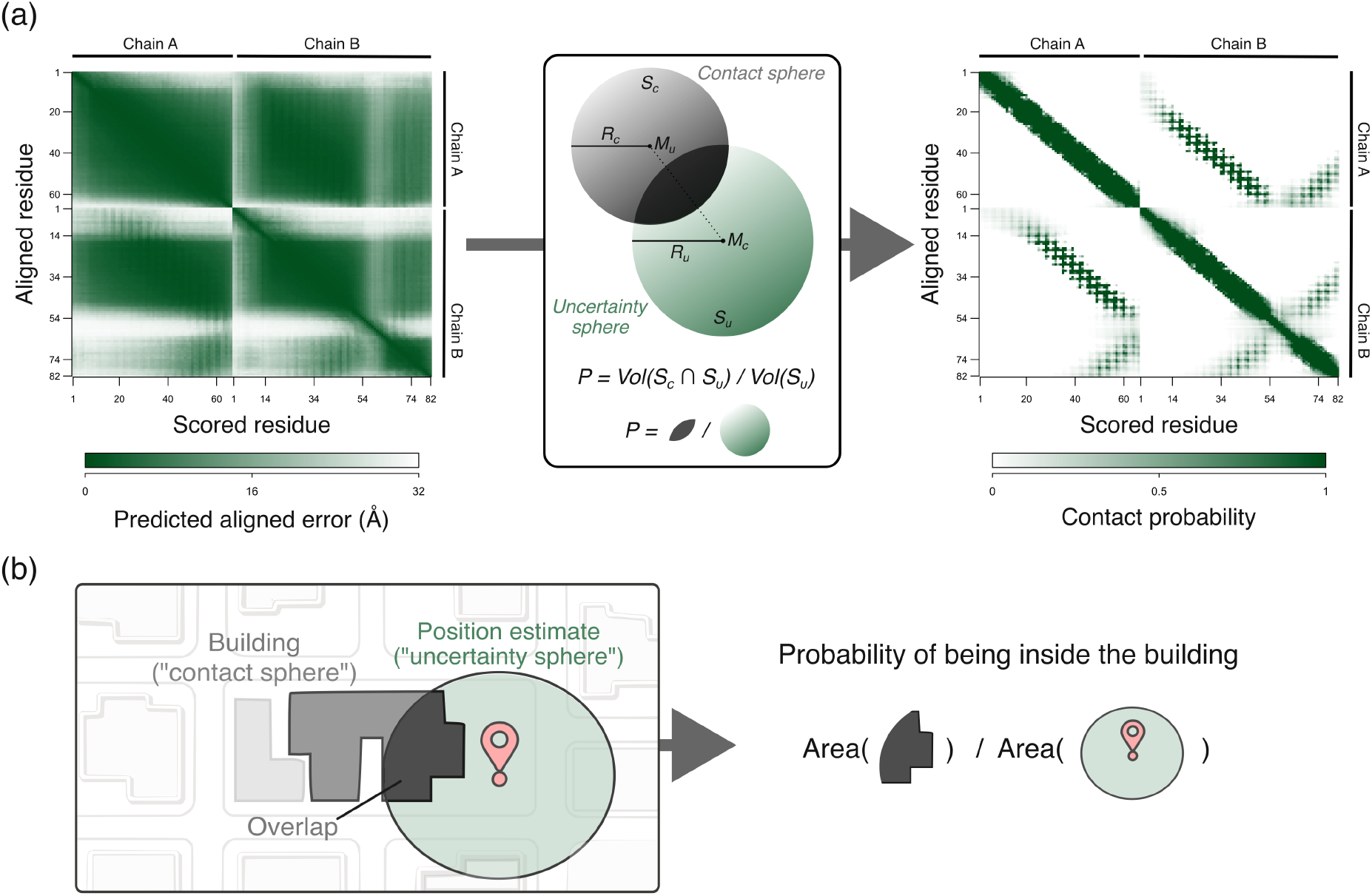
Overview of the derivation of Pinc. (**P**robability of **i**nterface **n**ative **c**ontacts). (**a**) *Left:* Each element in the PAE matrix represents the expected positional error of the scored residue when the aligned residue is correctly positioned. *Middle:* Consider two residues *C* (scored) and *U* (aligned), represented by their mass centres *M*_c_ and *M*_u_, respectively. The scored residue *C* is surrounded by an uncertainty sphere *S*_u_ with radius *R*_u_ corresponding to its PAE value (PAE[*U, C*]). The aligned residue *U* is surrounded by a fixed contact sphere *S*_c_ with radius *R*_c_. The contact probability is given by the volume of the intersection of *S*_c_ and *S*_u_ divided by the volume of the uncertainty sphere *S*_u_. *Right:* The resulting contact probability matrix is calculated using the predicted structure geometry. The Pinc score is computed as the mean contact probability for all interchain residue pairs with mass centres within *R*_c_ distance of each other. (**b**) The geometric model is analogous to GPS-based location estimation: the probability of being inside a nearby building is the fraction of the position uncertainty region that overlaps the building’s interior.

This model is analogous to GPS-based location estimation, illustrated in Figure 1b. A GPS receiver reports a position and an uncertainty radius, defining the region within which the true location is likely to lie. Consider a person standing near the entrance of a building. Depending on their actual position within this region, they may be inside or outside, and under the assumption that all positions are equally likely (isotropy) the probability of being inside is the fraction of the uncertainty region overlapping the building’s interior. A residue’s PAE value plays the same role by defining a sphere of plausible positions, where the contact probability is the fraction of this sphere that lies within contact distance of the aligned residue.

We extend this framework to multi-chain PAE matrices and use a centre-of-mass distance reprsentation computed over all heavy atoms within each residue. This choice reflects the observation that protein–protein interactions are predominantly mediated by side-chain contacts (Kortemme and Baker 2002; Talavera et al. 2011). Averaging over heavy atoms yields a distance measure that is less sensitive to local noise while retaining information about side-chain orientation. The Pinc score is then defined as the mean contact probability across all interchain residue pairs whose mass centres lie within the same contact radius used to derive the contact probabilities (fixed at 12 Å, see below).

Pinc is implemented as a standalone C program, with a Colab notebook available for convenient application to individual models. Given a structure file (PDB/mmCIF) and its matching PAE file (JSON), the program computes scores for all unique chain pairs by default. Optional flags allow outputting the full contact probability matrix and a list of non-zero interchain contact probabilities. Full documentation and source code are available at https://git.mpi-cbg.de/tothpetroczylab/Pinc.

### 2.2 Contact probabilities reflect the fraction of native contacts

To evaluate how well contact probabilities derived from our model (Figure 1) correspond to true interface contacts, we used two protein interaction benchmarks: one reproduced in-house (Dunbrack Jr 2025) and one analysed as provided (Genz et al. 2025). All models were generated without templates and correspond to dimers whose experimentally determined structures were deposited in the PDB after AlphaFold’s training cutoff date, with target sequences showing low homology to structures available before that date.

We computed interchain contact probabilities for the predicted models and grouped them into ten equidistant bins. For each bin, we calculated the fraction of interface contacts (Fnat) in the corresponding experimental structure. In both datasets, contact probabilities were well calibrated to the observed native contact fractions, with calibration curves lying close to the diagonal (Figure 2). Because calibration implies that each contact probability approximates the expected value of the corresponding native contact indicator, the Pinc score — their mean across the interface — is, by linearity of expectation, an approximately unbiased estimator of Fnat.

**Figure 2:**
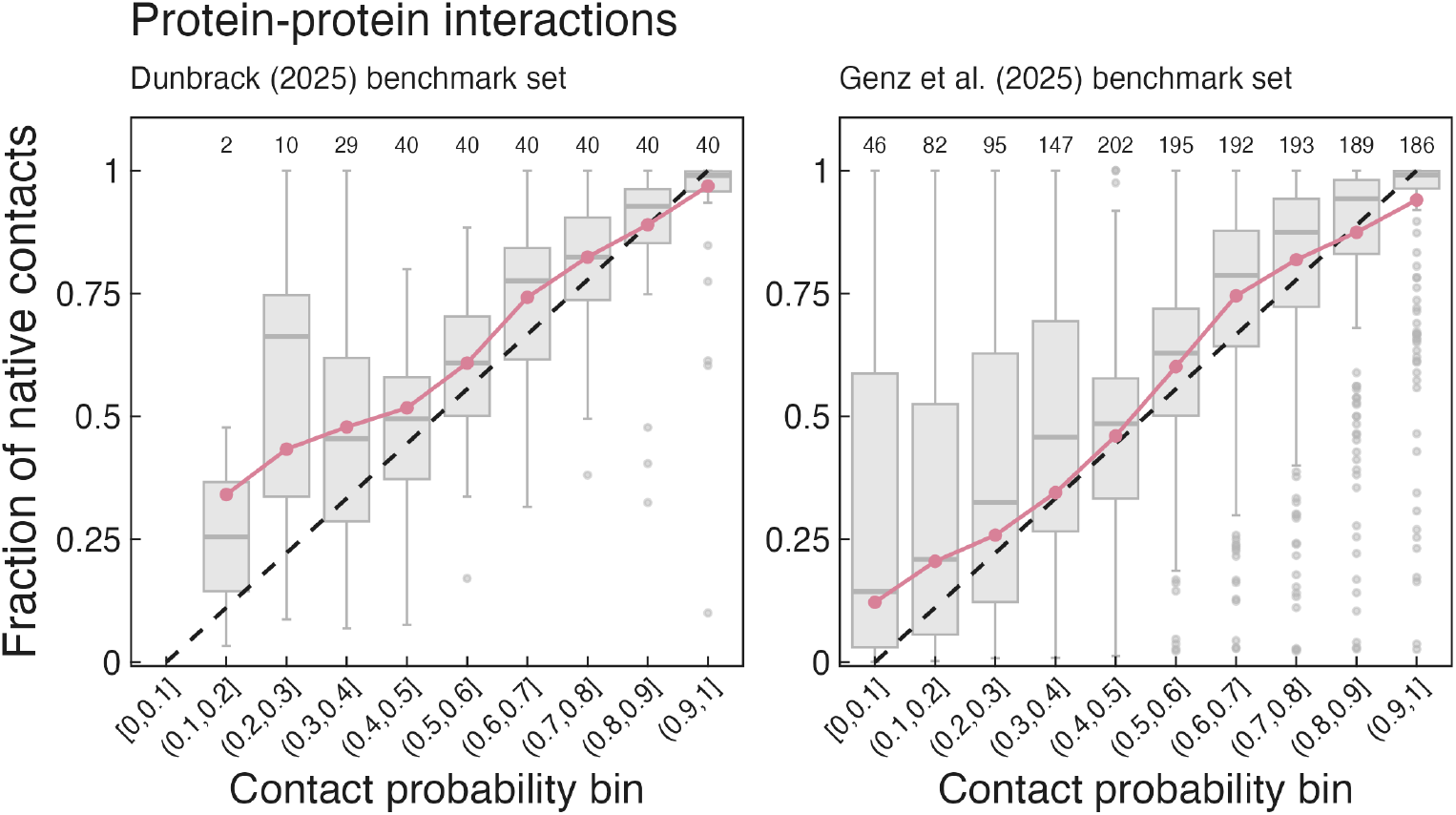
Residue contact probabilities are well-calibrated against experimental structures. Dunbrack (*left*) and Genz et al. (*right*) benchmark sets. Interchain residue–residue contact probabilities were computed from the AlphaFold PAE matrix using the geometric model (Figure 1) and grouped into ten equidistant probability bins. For each bin, the observed fraction of native interfacial contacts was calculated from the corresponding experimentally determined structures at a contact distance of 12 Å. The line and points show the global fraction of native contacts per bin, computed across all structures. Background boxplots summarise the structure-level distribution within each bin: boxes span the interquartile range, the middle line indicates the median, whiskers extend to 1.5× the IQR, and dots represent outliers. Note that the global fraction need not coincide with the mean of the structure-level distribution, as it is weighted by the number of contacts per structure rather than averaged across structures. The dashed diagonal indicates perfect calibration. Numbers along the top of each panel indicate the number of structures contributing to each bin.

To assess the sensitivity of Pinc to the choice of contact radius and residue distance representation, we evaluated seven models differing in contact sphere radius (8, 10, 12, and 15 Å), aggregation of interface halves (minimum or maximum Pinc between the two chains), and distance metric (centre-of-mass versus minimum heavy-atom distance). Calibration curves were broadly consistent across all distance thresholds for both benchmark sets (Figure S1), and the correspondence between Pinc and Fnat varied little between models, with differences falling within confidence intervals (Figure S2). Therefore, in line with the optimal distance range reported for the ipSAE (Dunbrack Jr 2025) and iLIS (Kim et al. 2026) metrics, we fixed the contact radius at 12 Å. Although Pinc is defined as the mean across the interface, the program also outputs chain-specific scores, which is practical for one-vs-all screens where the structural context of the fixed partner is invariant across runs.

Protein–nucleic acid complexes remain a challenging class of targets for AlphaFold, and many predicted models have incorrect interface topologies. We therefore evaluated Pinc in this difficult setting using a diverse benchmark of complexes deposited after AlphaFold’s training cutoff date (Peng et al. 2025) and predicted without templates. As expected from the limited accuracy of the underlying structural models, contact probabilities were less well calibrated to native contacts (Figure S3), with 30 of 59 complexes (51%) showing no overlap between predicted and native contacts. This reduced performance reflects a limitation of AlphaFold rather than of the scoring framework itself, because when the predicted interface topology is incorrect, non-native contacts contaminate the probability estimates. Consistent with this interpretation, restricting the analysis to complexes with higher native contact recovery (Fnat > 0.5) substantially improved calibration (Figure S3). Our program computes Pinc for all interacting chains, but we recommend treating Pinc scores for protein–nucleic acid interactions with caution, and we expect their utility to improve as AlphaFold models for protein–nucleic acid complexes become more accurate.

### 2.3 Pinc agrees with established interaction confidence scores

Using the contact probabilities derived above, we calculated Pinc scores for all predicted dimers in the Genz et al. dataset. We then compared Pinc to different interaction confidence scores, including pDockQ (Bryant et al. 2022), ipTM (Evans et al. 2021), iLIS (Kim et al. 2026), and ipSAE (Dunbrack Jr 2025). Pinc exhibited strong positive correlations with all other metrics (Figure 3a). The highest correlations were observed with ipTM, iLIS, and ipSAE (Spearman’s *ρ* = 0.96, *ρ* = 0.97, and *ρ* = 0.95, respectively).

**Figure 3:**
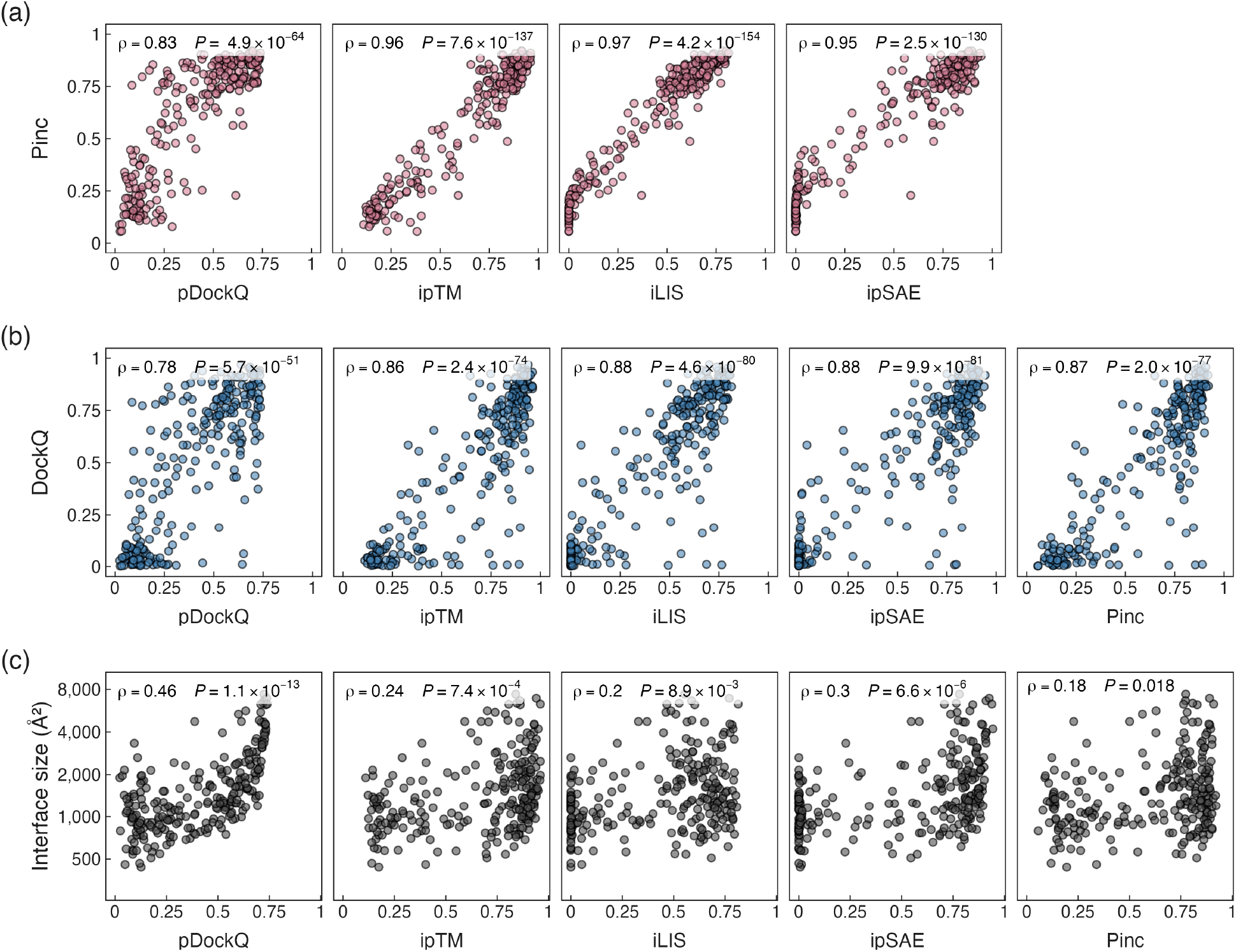
Pinc correlates with established interface confidence metrics. (**a**) Correlation of Pinc with commonly used AlphaFold interface confidence scores — pDockQ, ipTM, iLIS, and ipSAE — across the Genz et al. dimer benchmark set (*N* = 249). (**b**) Correlation of each confidence metric with DockQ, a composite measure of model quality based on native contacts, ligand RMSD, and interface RMSD relative to the experimental structure. (**c**) Correlation of each confidence metric with interface size, defined as the mean buried surface area across the two interface halves. All correlations are reported as Spearman’s *ρ* with Bonferroni-corrected *P*-values.

We next compared the correlation of confidence scores with the DockQ score, which quantifies the structural similarity between predicted and experimental structures by considering the fraction of native contacts, the root-mean-square deviation (RMSD) of the smaller chain, and that of the interface residues. Although DockQ does not account for potential conformational changes in the native dimer, it remains a suitable measure of structural similarity, conceptually independent from the TM-score. The results suggest high correlation between DockQ and Pinc (*ρ* = 0.87; Figure 3b), with the nominally highest correlation observed for ipSAE (*ρ* = 0.88).

Since the iLIS method demonstrated improved performance for small interfaces (Kim et al. 2026), we examined the relationship between confidence metrics and interface size, defined as buried surface area between the subunits (Figure 3c). The strongest correlation was observed for pDockQ (*ρ* = 0.46), consistent with its explicit dependence on the number of interface contacts. ipSAE, ipTM, and iLIS showed progressively weaker associations (*ρ* = 0.30, 0.24, and 0.20, respectively), with Pinc showing the weakest albeit significant correlation (*ρ* = 0.18, *P* = 0.018), suggesting that Pinc is the least influenced by interface size among the metrics evaluated.

### 2.4 Pinc is competitive with established metrics across independent benchmarks

We next evaluated the ability of Pinc and other confidence metrics — ipTM, ipSAE, and iLIS — to discriminate binders from non-binders across five benchmark sets: protein dimers (Dunbrack Jr 2025), protein multimers (Genz et al. 2025), de novo protein complexes (Overath et al. 2025), short linear motif (SLiM)-mediated protein dimers (Veinstein et al. 2025), and protein–nucleic acid complexes (Peng et al. 2025). In the SLiM-mediated protein dimers, we additionally included AlphaSLiM, a scoring metric developed in the benchmark study (Veinstein et al. 2025). Classification performance was assessed using the area under the receiver operating characteristic curve (AUROC) and the area under the precision–recall curve (AUPRC). Benchmark construction and class definitions are described in the Methods.

Performance across interaction classes broadly followed expectation: protein dimers and multimers were predicted with the highest accuracy, while SLiM-mediated and protein–nucleic acid complexes showed lower AUROC values (Figure 4a). All metrics exhibited intermediate performance on de novo protein complexes (Overath et al. 2025), which consist of computationally designed protein binders paired with their intended target proteins rather than naturally evolved interaction partners. Because benchmark labels in this dataset reflect whether a designed binder was experimentally observed to bind rather than the correctness of a known biological interaction, classification depends not only on the accuracy of the predicted interface but on additional factors influencing the binding experiment, such as binding affinity, expression, and solubility. These complexes also lack the evolutionary constraints and co-evolutionary signal typically available for natural protein complexes, representing a challenging but relevant use case for AlphaFold-based interface scoring. Performance on protein–nucleic acid complexes was slightly above chance for all metrics, in agreement with the calibration results discussed above.

**Figure 4:**
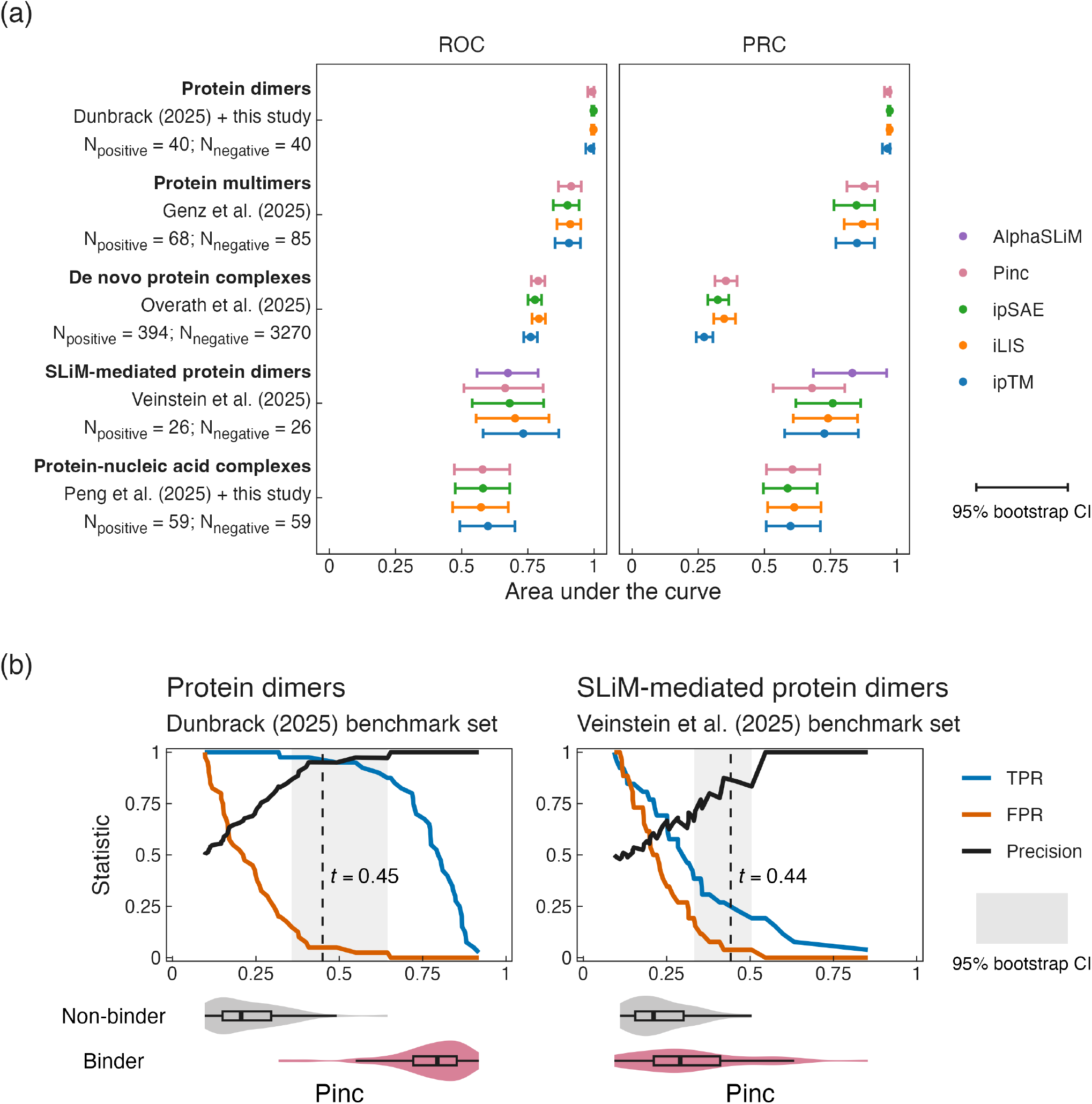
Performance of Pinc across diverse interaction classes. (**a**) Classification performance of Pinc, ipTM, iLIS, and ipSAE across five benchmark datasets spanning protein dimers, protein multimers, de novo protein complexes, SLiM-mediated protein dimers, and protein–nucleic acid complexes; axis labels annotated “+ this study” indicate datasets for which predictions, negative controls, or both were newly generated for this work rather than taken as published (see Methods). Performance is measured by the area under the receiver operating characteristic curve (ROC) and the area under the precision–recall curve (PRC). Points and error bars show estimates and 95% bootstrap confidence intervals (*N* = 10, 000 resamples). AlphaSLiM is included for the Veinstein et al. dataset only. (**b**) Classification threshold analysis for Pinc on the Dunbrack protein dimer and Veinstein et al. SLiM-mediated dimer benchmark sets. True positive rate (TPR), false positive rate (FPR), and precision are shown as a function of the Pinc threshold. The dashed vertical line indicates the optimal threshold *t* at 5% FPR, with the shaded region denoting the 95% bootstrap confidence interval. Violin plots show the distribution of Pinc scores for binders and non-binders in each dataset.

Pairwise bootstrap tests of the AUROC difference between Pinc and each comparator metric yielded no significant differences after Bonferroni correction in 14 of 16 comparisons (Figure 4a). The two exceptions, both in the Overath et al. de novo binder dataset, showed Pinc to perform significantly better than ipTM (*P* = 3.2 ×10^−3^) and ipSAE (*P* = 0.013). This is consistent with the interpretation that the geometric model underlying Pinc captures the information relevant for interaction confidence while making fewer assumptions and avoiding unnecessary complexity. We note, however, that the present benchmarks were not designed to discriminate between metrics, and larger and more carefully designed datasets with greater statistical power will be needed to determine whether any metric offers a meaningful advantage in specific interaction classes.

To facilitate the use of Pinc in interaction screens, we determined classification thresholds based on a target false positive rate of 5%, as this directly controls the fraction of non-binders incorrectly classified as binders, the error most consequential in a screening context. Using the protein dimer benchmark, we obtained a threshold of *t* = 0.45 (95% CI: 0.36–0.64), and using the SLiM-mediated dimer benchmark, *t* = 0.44 (95% CI: 0.33–0.50) (Figure 4b). As both benchmark sets comprise complexes with no close homologues in the PDB, these thresholds are expected to generalise well to screening scenarios where the vast majority of targets lack experimentally resolved structures. Both confidence intervals include 0.5, the natural midpoint of the Pinc scale, and we suggest *t* = 0.45 for general interaction screens, or the upper bounds of the intervals in more conservative settings.

### 2.5 Contact probabilities prioritise interface hotspot residues

Residues at protein–protein interfaces are commonly classified into three structural regions based on their relative solvent-accessible surface area in the unbound and bound states: support (partially buried in both states), core (exposed when unbound and buried upon binding), and rim (partially exposed in both states) (Levy 2010). Since contact probabilities are derived from the PAE matrix, which reflects AlphaFold’s confidence in the relative positioning of residue pairs, we expected them to vary between these regions according to how strongly the sites are constrained by the interface.

In the Genz et al. benchmark set, buried surface area followed the order support < rim < core, whereas residue contact probabilities followed the order rim < support < core (Figure 5a). Core residues are deeply buried at the interface, highly constrained by intermolecular contacts, and are the most evolutionarily conserved interface residues (Levy 2010). Together, these structural and evolutionary properties favour high-confidence predictions by AlphaFold and, consequently, high contact probabilities. Rim residues remain relatively exposed after binding and are less evolutionarily conserved, leading to lower contact probabilities. Support residues contribute little buried surface area but share the protein interior-like environment of core residues in both the bound and unbound states, yielding contact probabilities that exceed those of rim residues. This distinction is notable because the impact of mutations on binding free energy precisely follows this same ordering (rim < support < core) (David and Sternberg 2015; Petukh et al. 2015), which is recapitulated in the depletion of disease-associated variants at the rim (Livesey and Marsh 2022). These findings suggest that contact probability reflects local structural constraint rather than interfacial burial alone, offering a residue-level means of prioritising mutationally sensitive interface positions.

**Figure 5:**
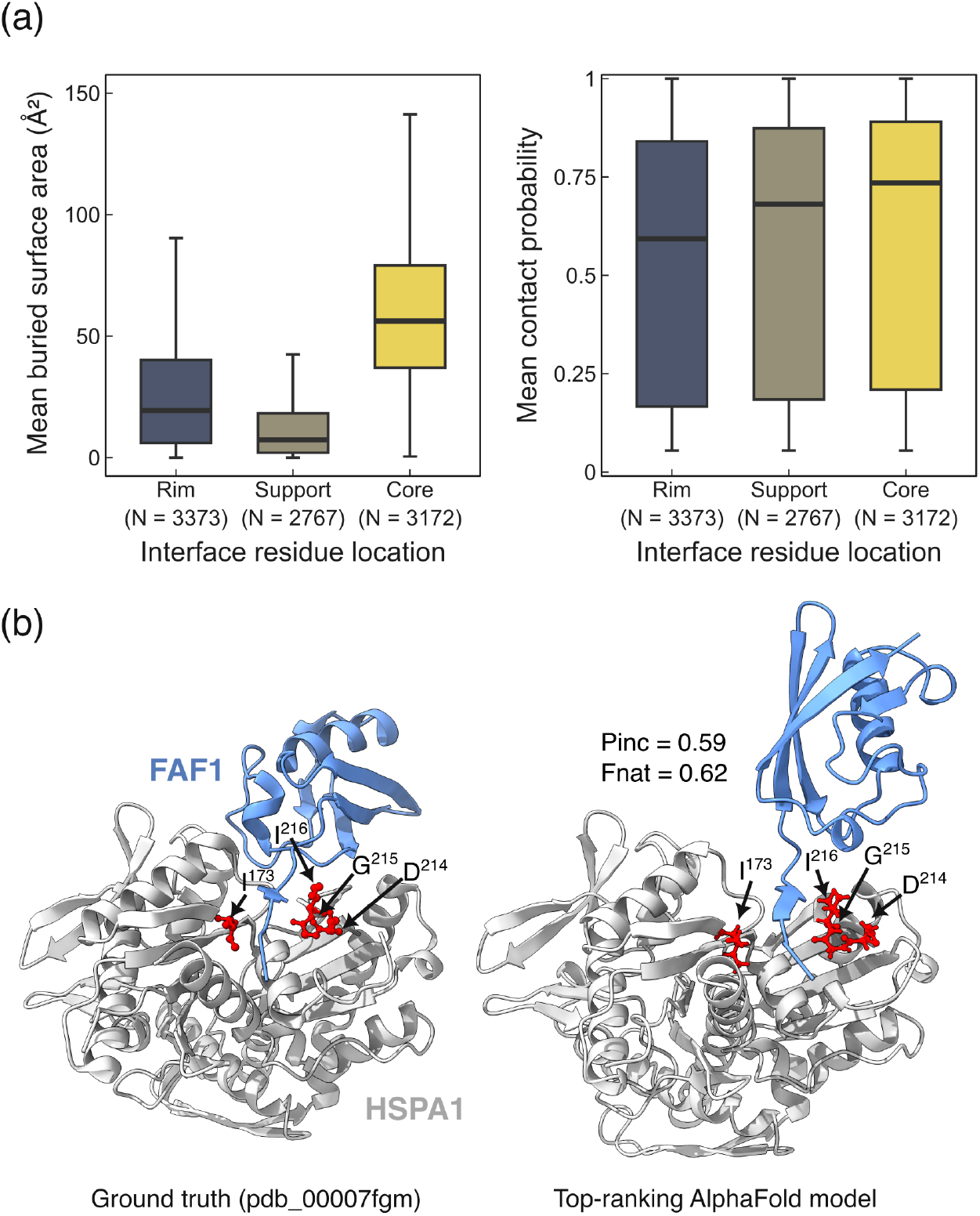
Residue contact probabilities identify interface hotspots. (**a**) Mean buried surface area (*left*) and mean residue contact probability (*right*) across interface residue locations in the Genz et al. benchmark set. Boxes span the interquartile range, the middle line indicates the median, and whiskers extend to 1.5× the IQR. *N* denotes the number of residues in each group. (**b**) *Left:* Experimentally determined structure of FAF1(L-UBL1):HSPA1. *Right:* Top-ranking AlphaFold model for the FAF1(L-UBL1):HSPA1 interaction. Although the L-UBL1 domain of FAF1 is positioned differently in the model, binding of the *β*-structured linker is nonetheless correctly predicted. Residues with high contact probabilities (*P* > 0.95) are shown as orange spheres, marking a conserved hotspot on the HSPA1 surface.

As an illustration of how residue contact probabilities can be used to identify interface hotspots, we consider the interaction between FAF1(L-UBL1) and HSPA1 (Figure 5b). FAF1 acts as a molecular scaffold that recruits IQGAP1 to the HSPA1 chaperone; this complex suppresses RhoA signalling to stabilise adherens junctions and maintain tissue integrity (Song et al. 2022). Although the relative orientation of the L-UBL1 domain differs between the experimental structure and the top-ranking AlphaFold model, the binding mode of the *β*-structured linker is nonetheless correctly predicted. High residue contact probabilities (*P* > 0.95) are assigned to a small set of residues lining the HSPA1 surface, defining a compact interaction hotspot in the experimental structure that can be identified despite global modelling inaccuracies. To facilitate such analyses, our program reports both interface-level Pinc scores and residue-residue contact probabilities as output.

## 3 Discussion

We introduce Pinc, an interaction confidence score that converts AlphaFold PAE values into contact probabilities, providing an estimate of the fraction of native interfacial contacts. Pinc shows consistent calibration to contact fractions observed in experimental structures and strong agreement with existing confidence metrics. This relationship allows interpretation of model confidence in biological terms: a score of 0.8 can be read as predicting that around 80% of the native contacts are present in models with that score. Unlike metrics aimed at approximating structural similarity measures, Pinc offers an intuitive probabilistic framework for assessing predicted protein–protein interfaces.

A key advantage of Pinc is that it depends on a single empirically fixed parameter, the contact radius, and requires only the predicted structure and its PAE matrix. This contrasts with actifpTM (Varga et al. 2025), which depends on distograms that are not typically retained during inference, making post hoc computation impractical. Where ipSAE (Dunbrack Jr 2025) and iLIS (Kim et al. 2026) impose discrete distance and PAE cutoffs that require careful calibration, Pinc instead propagates positional uncertainty continuously into a contact probability, avoiding the need to threshold the PAE. Unlike ipTM, Pinc is by construction independent of total chain length, as only the residues contributing to the interface enter the calculation. Consistent with this design, Pinc showed the weakest dependence on interface size of all metrics evaluated, suggesting it is comparatively more sensitive to the quality of individual interfacial contacts. This is further supported by our finding that contact probabilities reflect local structural constraint rather than interfacial burial alone.

Across five independent benchmark sets spanning diverse interaction classes, Pinc’s classification performance was statistically indistinguishable from established metrics in most pairwise comparisons, with exceptions confined to the de novo protein complex benchmark, where Pinc outperformed ipTM and ipSAE (Figure 4a). We interpret this finding cautiously, as the present benchmarks were not designed to discriminate between scoring methods, and larger, carefully designed datasets with greater statistical power will be needed to determine whether any metric offers a meaningful advantage in specific interaction classes. Within this limitation, however, performance parity combined with a simpler and more interpretable formulation provides a practical argument for Pinc as a first-line metric, particularly for interaction classes or experimental contexts where established metrics have not been independently validated. For binary interaction screens, a Pinc threshold of 0.45 provides a practical cutoff applicable to both stable and motif-mediated complexes (Figure 4b), above which fewer than 5% of predicted interactions are expected to be false positives.

The performance of all metrics, including Pinc, was considerably lower for protein–nucleic acid interactions than for protein–protein interactions, consistent with previous observations (Elofsson 2026; Peng et al. 2025; Xu et al. 2025). We attribute this to the current limitations of AlphaFold in accurately predicting the topology of protein–nucleic acid interfaces, rather than to a fundamental limitation of the contact probability framework. When the predicted interface is broadly correct, as in the subset of complexes with higher native contact recovery, calibration improved substantially. As AlphaFold’s accuracy for this interaction class continues to develop, we expect the utility of Pinc, and of contact probability-based scoring more generally, to grow correspondingly.

Several caveats apply when interpreting Pinc scores. The contact radius is fixed in the implementation to ensure comparability across datasets, and was selected empirically rather than on structural or physical grounds. Pinc is not a direct substitute for metrics such as ipTM, which imposes more explicit spatial constraints in principle; yet correlates with DockQ to a similar extent, indicating that a contact-based probabilistic formulation captures much of the relevant structural information necessary to evaluate interaction confidence. We therefore propose Pinc as a complementary metric: one that makes fewer assumptions than existing methods, is straightforward to compute and interpret, and whose probabilistic nature is preserved at the residue level, making it well suited for identifying interface hotspots and prioritising positions for mutational studies.

## 4 Methods

### 4.1 Structural datasets

#### Protein dimers (Dunbrack dataset)

Protein Data Bank [PDB; Berman et al. (2000)] identifiers for the 40 heterodimers were acquired from the preprint (v2) of Dunbrack Jr (2025). The dataset was constructed to minimise homology to structures available before the AlphaFold-Multimer v2.3 training cutoff of 30 September 2021: all entries share at most 40% sequence identity with any chain deposited in the PDB before that date. Each entry contains exactly two unique sequences forming a biological assembly, with each chain comprising at least 12 residues. Chain sequences were extracted from the PDB SEQRES records. AlphaFold2-Multimer predictions were generated with ColabFold (Kim et al. 2025; Mirdita et al. 2022) using LocalColabFold v1.5.5 on a single NVIDIA RTX A6000 GPU (40 GB RAM), with multiple sequence alignments constructed using the mmseqs2_uniref_env_envpair setting. Five models were generated across five random seeds (42–46), each using three recycles, yielding 25 models per dimer. For a negative set, we constructed 40 non-native heterodimer pairs by randomly reassigning sequences across the 40 PDB entries, predicted under identical settings.

#### Protein dimers (Genz et al. dataset)

We used template-free AlphaFold2-Multimer predictions generated with ColabFold and made publicly available at https://gitlab.com/topf-lab/c2qscore (Genz et al. 2025). The dimer benchmark comprises PDB entries released between 1 October 2021 and 11 July 2025 at ≤ 3.0 Å resolution, grouped at 30% sequence identity to ensure non-redundancy, and filtered to retain heterodimeric complexes with at least three residues per chain in contact at a 5 Å heavy-atom cutoff, chains of at least 50 residues, no antibody or nanobody components, and a biological assembly consistent with the asymmetric unit. After processing, 249 predicted dimers were retained for calibration and correlation analyses throughout this study; ten overlap with the Dunbrack dataset.

#### Protein multimers

For the classification benchmark only, we used the large assembly dataset of Genz et al. (2025), comprising dimers derived from higher-order assemblies resolved by cryo-EM at ≤ 2.3 Å, deposited after 1 September 2023, and curated under the same criteria as the dimer benchmark. After processing, 153 dimers were retained. Predictions were labelled using CAPRI quality criteria: models with DockQ ≥ 0.23 were considered acceptable or better and served as the positive class, while those with DockQ < 0.23 formed the negative class. This framing does not represent true non-binders well, as the negative class consists of incorrectly modelled rather than non-interacting pairs, and should be interpreted as a proxy for model quality discrimination rather than interaction prediction. Since all metrics are evaluated under identical conditions, the dataset remains appropriate for comparing relative classification performance, and we do not derive or recommend classification thresholds from it.

#### De novo protein complexes

We used the compiled dataset of de novo miniprotein binders and associated AlphaFold2 predictions made publicly available by Overath et al. (2025) at https://doi.org/10.5281/zenodo.15722219. The dataset comprises 3,766 unique binder designs targeting 15 distinct proteins, of which 436 (11.6%) were confirmed as in vitro binders and served as the positive class; the remaining designs formed the negative class. Binding labels were derived from experimental assays that varied in format and affinity threshold across source studies. After processing, all scores evaluated in this study could be computed for 3,664 complexes.

#### SLiM-mediated protein dimers

We used the noPDB_dataset subset of the ELM-derived benchmark reported in Veinstein et al. (2025), comprising 26 SLiM-mediated protein interactions with no corresponding structural information in the PDB, which served as the positive class. A matched negative set of 26 non-native pairs was constructed by the original authors by randomly shuffling SLiM–target combinations from the full benchmark, excluding pairs in which the SLiM matched any regular expression associated with the new target. AlphaFold predictions and confidence scores were used as provided by the authors and are available via the associated repositories.

#### Protein–nucleic acid complexes

We reconstructed the Benchmark-IX protein–nucleic acid dataset from the 59 complexes reported in Peng et al. (2025). The dataset comprises structures deposited after 1 May 2023, ensuring all entries post-date the AlphaFold3 training cutoff. Homology to the training set was controlled by excluding complexes where all protein chains shared > 40% sequence identity, or all nucleic acid chains showed significant similarity (BLASTN *E*-value < 10), with any entity in the AlphaFold3 training data. Structural redundancy was further reduced by clustering at a TM-score threshold of 0.5 and retaining the largest representative per cluster. Additional filters required a total residue count of < 1000, at least 10 nucleotides per chain, and at least 80% structural coverage of both components. Chain sequences were extracted from the PDB SEQRES records. Predictions were generated via the AlphaFold3 server (Abramson et al. 2024) with templates excluded, run on two consecutive days starting on 13 February 2026 using the default random seed. For a negative set, we constructed 59 non-native protein–nucleic acid pairs by selecting, for each complex, the protein chain at the largest protein–nucleic acid interface, and pairing it with a nucleic acid chain from a different complex, matched by nucleic acid type and selected to maximise sequence dissimilarity to the native partner while penalising large differences in chain length. These pairs were predicted under identical settings, run on two consecutive days starting on 10 June 2026 with the default random seed.

### 4.2 Residue mapping

Residue correspondence between experimental structures and AlphaFold-predicted models was established by pairwise sequence alignment using MUSCLE v5.3 (Edgar 2022). For each complex, the sequences extracted from the predicted and experimental structure coordinates were aligned per chain, and the resulting alignments were used to construct a residue index map between the two structures. Only residues present in both structures were retained.

### 4.3 Calculation of structural metrics

Confidence scores ipTM, iLIS, and ipSAE were computed using the lis.py script (Kim et al. 2024) with default settings (https://github.com/flyark/AFM-LIS/blob/main/lis.py). DockQ and pDockQ scores for the Genz et al. dataset were computed by the original authors and obtained from the associated GitLab repository. Interface size was defined as the mean buried surface area across the two interface halves and computed using FreeSASA v2.1.2 (Mitternacht 2016) with default parameters.

Where multiple models were available, all metrics were averaged across models to reduce potential effects of model ranking: across the top ten models for the Dunbrack dataset, following Dunbrack Jr (2025); across all five models for the Genz et al. and Overath et al. datasets. For the Veinstein et al. dataset, scores from the top-ranking model were used to retain comparability with the AlphaSLiM score. For the Peng et al. dataset, given the generally poor agreement between predicted and native contacts in this protein–nucleic acid setting, scores from the top-ranking model were used.

Interface residues were classified into core, rim, and support categories following the RSA-based scheme of Levy (2010). RSA was defined as the solvent-accessible surface area of a residue normalised by its maximum accessible surface area based on Gly-X-Gly tripeptide reference values (Miller et al. 1987). Residues with positive buried surface area upon complex formation were assigned to one of three categories: core (RSA ≥ 0.25 unbound, < 0.25 bound), rim (RSA ≥ 0.25 in both states), or support (RSA < 0.25 in both states).

### 4.4 Bootstrap procedure

All confidence intervals were computed from 10,000 bootstrap replicates, with the 2.5th and 97.5th percentiles as the interval bounds. For classification metrics, resampling was stratified by class to preserve the binder-to-non-binder ratio. Pairwise differences in AUROC between metrics were assessed using a paired, stratified bootstrap difference test, with two-sided *P* -values derived from the bootstrap distribution using continuity correction (Davison and Hinkley 1997); multiple comparisons were adjusted with the Bonferroni method.

### 4.5 Statistical analysis

All data visualisation and statistical analyses were performed in R v4.5.3 (R Core Team 2026).

### 4.6 Molecular visualisation

Structural figures were created in ChimeraX v1.10.1 (Meng et al. 2023).

## Acknowledgements

We thank the Computer Services and Scientific Computing Facilities at MPI-CBG for their support, Anna Hadarovich for testing the original R script and Colab notebook, and Swantje Lenz for helpful discussions on the manuscript.

## Data availability statement

The C program and a Colab notebook to calculate the Pinc score are available at https://git.mpi-cbg.de/tothpetroczylab/Pinc. Associated data and code to reproduce the results are available at https://doi.org/10.17617/3.SVPIUW.

