## Supplementary figures for "Pinc: a simple probabilistic AlphaFold interaction score"

### Supplementary material for
### Pinc: a simple probabilistic AlphaFold interaction score

Mihaly Badonyi[1,2,3†] and Agnes Toth-Petroczy[1,3,4*]

[1]Max Planck Institute of Molecular Cell Biology and Genetics, Pfotenhauerstraße 108, 01307, Dresden, Germany
[2]Max Planck Institute for the Physics of Complex Systems, Nöthnitzer Straße 38, 01187 Dresden, Germany
[3]Center for Systems Biology Dresden, Pfotenhauerstraße 108, 01307, Dresden, Germany
[4]Cluster of Excellence Physics of Life, TU Dresden, Arnoldstraße 18, 01062, Dresden, Germany

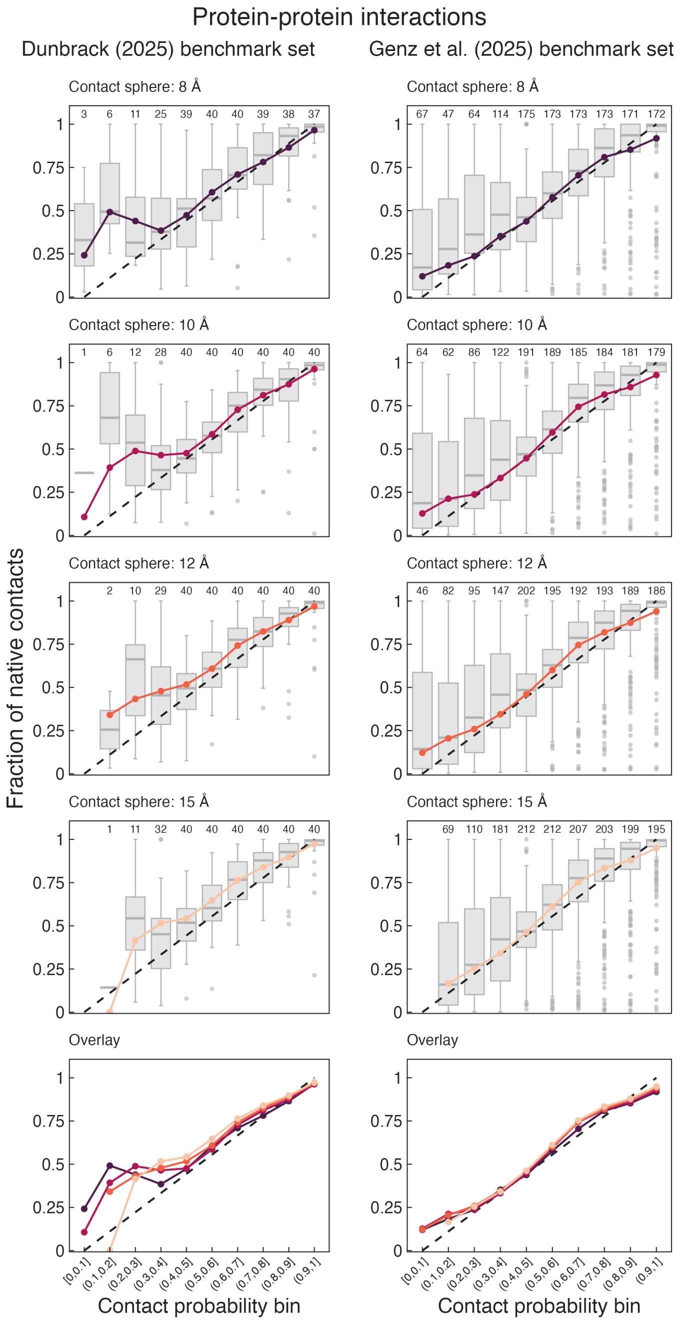


**Figure S1: Calibration curves across contact distance thresholds for protein–protein interactions.** Dunbrack (*left*) and Genz et al. (*right*) benchmark sets. Interchain residue–residue contact probabilities were computed from the AlphaFold PAE matrix using the geometric model and grouped into ten equidistant probability bins, as in Figure 2. Each row corresponds to a different contact distance threshold (8, 10, 12, and 15 Å), with the bottom row showing all thresholds overlaid. For each bin, the observed fraction of native interfacial contacts was calculated from the corresponding experimentally determined structures. The line and points show the global fraction of native contacts per bin, computed across all structures. Background boxplots summarise the structure-level distribution within each bin: boxes span the interquartile range, the middle line indicates the median, whiskers extend to 1.5× the IQR, and dots represent outliers. The dashed diagonal indicates perfect calibration. Numbers along the top of each panel indicate the number of structures contributing to each bin.

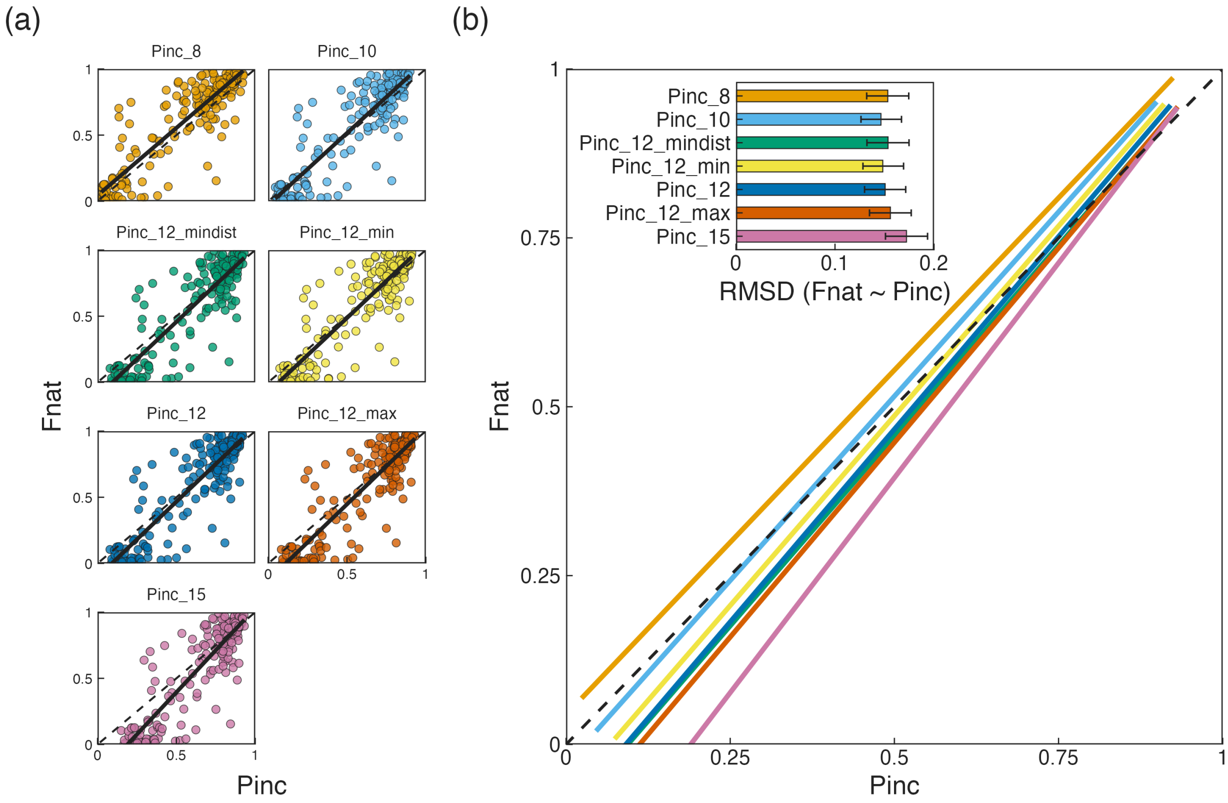


**Figure S2: Correspondence between Pinc and Fnat across parameter settings.** (**a**) Scatterplots of Pinc against the fraction of native contacts (Fnat) for seven Pinc models differing in contact distance threshold and distance representation, evaluated on the Genz et al. dimer benchmark set. Each panel shows a linear regression line (black) and the identity line (dashed diagonal). Points are coloured by model. Only structures with Fnat $> 0$ are included ($N = 223$). (**b**) Overlaid regression lines for all seven models, highlighting differences in slope across parameter settings. The inset shows RMSD between Pinc and Fnat for each model, with 95% bootstrap confidence intervals ($N = 10,000$ resamples). Differences across models are small and largely within confidence intervals, supporting the robustness of Pinc to the choice of distance threshold. Model suffixes denote parameter settings: numbers indicate the contact sphere radius in Å; `_min` and `_max` denote the minimum and maximum Pinc across the two interface halves, respectively; and `_mindist` indicates that minimum heavy-atom distance was used in place of centre-of-mass distance when computing the residue distance matrix.

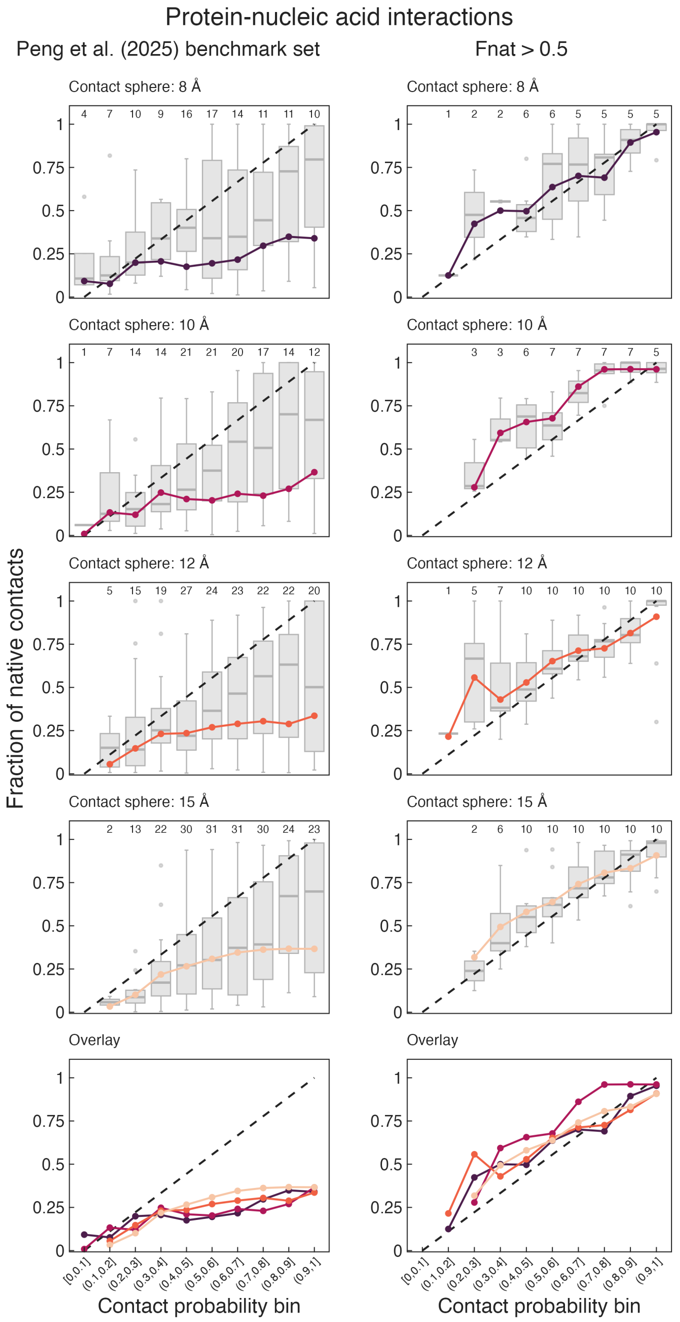


**Figure S3: Calibration curves across contact distance thresholds for protein–nucleic acid interactions.** Peng et al. benchmark set, full dataset (*left*) and restricted to complexes where native contacts comprise more than 50% of all predicted contacts (Fnat $> 0.5$; *right*). Interchain residue–residue contact probabilities were computed from the AlphaFold PAE matrix using the geometric model and grouped into ten equidistant probability bins, as in Figure 2. Each row corresponds to a different contact distance threshold (8, 10, 12, and 15 Å), with the bottom row showing all thresholds overlaid. For each bin, the observed fraction of native interfacial contacts was calculated from the corresponding experimentally determined structures. The line and points show the global fraction of native contacts per bin, computed across all structures. Background boxplots summarise the structure-level distribution within each bin: boxes span the interquartile range, the middle line indicates the median, whiskers extend to 1.5× the IQR, and dots represent outliers. The dashed diagonal indicates perfect calibration. Numbers along the top of each panel indicate the number of structures contributing to each bin.
